# Polyolefin colonization and partial degradation by *Gordonia* sp., and *Arthrobacter* sp. isolated from wetlands and compost

**DOI:** 10.1101/2024.12.20.629628

**Authors:** Scott A. Coughlin, Adam McFall, Stephen A. Kelly, Julianne Megaw

**Affiliations:** School of Biological Sciences, Queen’s University Belfast, 19 Chlorine Gardens, Belfast, BT9 5DL, Northern Ireland; School of Pharmacy, Queen’s University Belfast, 97 Lisburn Road, Belfast, BT9 7BL, Northern Ireland; School of Medicine and Centre for Interventions in Infection, Inflammation, and Immunity (4i), University of Limerick, Limerick, Ireland

## Abstract

Plastics and microplastics constitute an ever-increasing pollutant in the biosphere. There is clear evidence that environmental microbes possess enzyme systems and metabolic apparatus to degrade natural high molecular weight polymers, which substantially overlap with those involved in the biodegradation of plastics. This study investigated the presence and activities of plastic-degrading microbes in environments with a high abundance of plant-derived polymers, including cellulosic, chitinous, or lignin-derived compounds. Microbes were enriched in a minimal medium, with low-density polyethylene (LDPE), polypropylene (PP), poly(ethylene terephthalate) (PET), or polystyrene (PS) as the sole carbon source, resulting in 12 bacterial isolates. Plastic mass loss and cell viability were measured over a 28-day incubation period, in addition to assessment of biofilm formation. *Gordonia* sp. iso11 displayed mass loss of PP of up to 22.8% and formed a viable dense biofilm (10^9^ CFU cm^-2^) on a PP film. A putative alkane-1-monoxygenase (AlkB) was identified from genome sequence analysis, which aligned with a known LDPE-degrading enzyme. Furthermore, reductions in the contact angle of medium supernatant from *Gordonia* sp. iso11 provided evidence of biosurfactant production which may enhance the bioavailability of the synthetic plastic. To our knowledge, this is the first demonstration of PP degradation by *Gordonia*.

## 1. Introduction

It is estimated that by 2040, global environmental plastic pollution will weigh a total of 1.3 billion tonnes [1], with highly recalcitrant plastic polymers becoming ubiquitous across virtually all biomes on Earth. Polyethylene (PE) and polypropylene (PP), and their soluble leachates, pose a threat to biodiversity, evidenced by growth prohibition and community flux of the phytoplankton at the base of the food web [2, 3]. Ingestion of microplastics in animals and zooplankton contribute to their global dispersal over prolonged periods of time [4-8]. This poses a risk to human food chains [9, 10]. Microplastic-induced fluxes in the macroecology of Earth’s biosphere have also been predicted [11]. Therefore, there is urgent need to devise methods for remediation of plastic-contaminated sites and waters. Similar to alkanes and other hydrocarbons, plastics are among the most energy-dense carbon sinks available to microbes [12]. Therefore, increasing attention has been drawn to identifying microbial and enzymatic systems with the ability to depolymerize plastics [13, 14]. Indeed, culture-based approaches to isolate plastic-degrading microbes have yielded a completely novel, and previously uncultured polyethylene terephthalate (PET)-degrading bacterium, *Ideonella sakaiensis* 201 F6, which uses a two-enzyme system to degrade, assimilate, and mineralize PET [15]. Furthermore, mutagenic approaches have enhanced the polymerization rates of PET by PETase from *I. sakaiensis* six-fold in a cell-free system [16, 17]. Whilst rapid rates of microbial PET mineralization have been observed, polypropylene (PP) is typically evidenced to be more recalcitrant towards microbial biodegradation; PP is more hydrophobic than PET [18] and studies involving a range of synthetic plastic often observe slower rates of degradation for PP [19, 20].

Some culture-based approaches have focused on the isolation of microorganisms from plastic-contaminated sites, which may be enriched for species that can metabolize such substrates. These have been successful in yielding microbes which rapidly degrade plastic. For example, a study in a plastic-contaminated Malaysian mangrove swamp resulted in the culture of two *Bacillus* spp., each capable of degrading PP, PE, PET, and polystyrene (PS) with varying rates [19]. In another study which sampled petroleum contaminated beach soil, *Pseudomonas aeruginosa* E7 was isolated, which degrades PE under aerobic conditions [21]. Many such studies utilized environments with long-term plastic and/or hydrocarbon pollution, providing an excellent rationale to screen for plastic-degrading microbes [22]. Indeed, a simulation of the conditions of the Deepwater Horizon oil spill, which occurred in the Gulf of Mexico in 2010, found very rapid enrichment of microbes capable of degrading hydrocarbons present in crude oil, with differential expression of hydrocarbon degrading enzymes [23]. Given the similarities of the hydrocarbon constituents of crude oil and synthetic plastic (which is often derived from crude oil), similar enrichment may occur in plastic-polluted environments.

Whilst many culture-based isolation studies have been carried out in environments which are extremely plastic-polluted (e.g. landfill sites), few studies have considered the potential of environments rich in plant polymers to harbor plastic-degrading microbes, despite many of the enzyme systems and metabolic pathways responsible for the biological breakdown of plastic are also those used in the breakdown of plant polymers [24].

We characterized *Gordonia* sp. iso11 (isolated from a wetland peat bog) in terms of its ability to secrete surfactants, form biofilms, and grow under nutrient stress. Bacteria from the genus *Gordonia* are pertinent in bioremediation of hydrocarbons; they display a high degree of metabolic plasticity, evidenced by the wide range of carbon substrates they can utilize as an energy source [25-28]. Critically, the microbes utilize enzyme systems, such as monooxygenases or P450 cytochromes, to convert long-chained hydrocarbons into alcohols, which are available for further chemical alteration by enzymatic catalysis to ultimately yield acetyl-CoA derivatives (from by β-oxidation of fatty acids), from which energy is derived in the TCA cycle.

This study enhances our understanding of the approaches which may be used to isolate novel plastic degraders, where a functional approach to bioprospecting is carried out across wetland environments.

## 2. Materials and methods

### 2.1. Plastics

Plastics used in the screening experiments were in the form of high-purity pellets, free from additives or other soluble components, sourced from Merck (Darmstadt, DE). The plastics used were PP (CAS 9003-07-0), LDPE (CAS 9002-88-4), PET (CAS 25038-59-9), and PS (CAS 9003-53-6).

Plastic mass loss assays utilized high purity plastic films, with the polymers biaxially orientated, free from, sourced from Goodfellow Cambridge Limited (Huntingdon, UK). The plastics used were PP (0.008 mm thickness), LDPE (0.2 mm thickness), PET (0.006 mm thickness), and PS (0.05 mm thickness).

### 2.2. Screening and enrichment of plastic-degrading activity of compost and peat bog microbes

Various microbial habitats which harbor insoluble polymers were selected. The samples collected were peat, lake sediment, reed-bed mud, and domestic compost. Samples were collected and placed into 50 ml sterile screw-top plastic tubes. Tubes were placed on ice during transport and stored at 4 °C for short term storage (up to 1 day).

From the wetland (peat bog) and compost samples, 5 g were suspended by vortex mixing in a 50 mL conical tube containing a 25 mL solution of 5% w/v NaCl. Sediment was left to settle, after which 1 ml of the suspension was used to inoculate 100 mL Erlenmeyer flasks containing 25 mL carbon-free Czapek-Dox medium (NaNO_3_ 3 g L^-1^; MgSO_4_ 0.5 g L^-1^; KCl 0.5 g L^-1^; K_2_PO_4_ 1.0 g L^-1^; FeSO_4_ 0.01 g L^-1^), supplemented with 2.5 g plastic pellets (PE, PP, PET, and PS), sterilized by soaking with 70% v/v ethanol and dried by evaporation in an aseptic environment. Flasks were incubated aerobically at 30 °C for 28 days, with shaking at 100 rpm. Growth in the flasks was determined by observing increases in the turbidity of the otherwise clear and colorless growth medium. Pellets were subsequently removed and used to inoculate Petri dishes containing Czapek-Dox agar, containing 0.5% w/v D-glucose as a soluble carbon source. These were incubated at 30 °C and any resulting bacterial colonies were isolated by streaking onto plates of the same medium.

### 2.3. Identification of isolates and phylogenetic analysis

The isolates were identified by 16S rRNA gene sequencing. Colony PCR was used to amplify the 16S rRNA gene of each isolate, using DreamTaq Green PCR mix (Thermo Scientific) and the universal bacterial primer pair 27F (5’-AGAGTTTGATCMTGGCTCAG-3’) and 1492R (5’-GGTTACCTTGTTACGACTT-3’). The presence and purity of PCR products was confirmed by agarose gel electrophoresis, after which the PCR product was purified using a GeneJET PCR purification kit (Thermo Scientific), with Sanger sequencing conducted by Eurofins Genomics (Cologne, DE). The sequences obtained were identified using NCBI BLAST.

A maximum likelihood tree was constructed using the 16S sequences obtained in this study, along with the 16S sequences of closely related strains and other plastic-degrading microbes, which were obtained from the NCBI GenBank database. This was carried out with bootstrapping (1000 replicates), with the Jukes-Cantor model was used to estimate the pairwise genetic distances. The PhyML workflow as used on NGPylogeny.fr [29], with tree annotation using ITOL [30].

### 2.4 Biodegradation activity of wetland and compost microbes

Degradation rates of plastic by the isolates were determined by gravimetric mass loss of high-purity plastic films (accurate to 0.02 mg). Films were cut into 20 x 40 mm pieces (corresponding to ∼6.3 mg), sterilized by soaking in 70% v/v ethanol and drying by evaporation in an aseptic environment. Cultures of *Gordonia* sp. iso11 and *Arthrobacter* sp. iso 16 were prepared by aerobic incubation in M9 broth (Na_2_HPO_4_ 6 g L^-1^, NaH_2_PO_4_ 3 g L^-1^, NH_4_Cl 1 g L^-1^, NaCl 0.5 g L^-1^, MgSO_4_ 0.12 g L^-1^) supplemented with 0.25% w/v D-glucose and 0.5% w/v yeast extract, at 37 °C until the cell density (OD_600_) reached > 0.6. From these cultures, 1 mL aliquots were removed and centrifuged, and the cell pellet was resuspended in carbon-free M9 medium. The resulting cell suspension was used to inoculate 100 mL Erlenmeyer flasks containing 20 mL carbon-free M9 medium (with 0.25% w/v casamino acids) and PP or PS film. The flasks were incubated at 37 °C for 28 days, with partial medium replenishment (10 mL) every 24 h. Following incubation, plastic films were removed and sonicated for 50 minutes in 25 mL sterile water in a 50 mL centrifuge tube, which was placed into an ultrasonic water bath (Branson M Series Ultrasonic Cleaning Bath, Branson Ultrasonics, CT, USA). Films were subsequently washed in 0.1% w/v sodium dodecyl sulfate (SDS) solution [31], rinsed in reverse-osmosis water, dried, and re-weighed. The procedure was carried out in triplicate, with uninoculated controls. Percentage mass change was given as

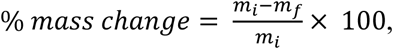

where *m_i_* is the initial mass of the polymer, and *m_f_* is the mass of the polymer at the end of the experiment.

### 2.5. Biodegradation activity in the presence of other carbon substrates

Biodegradation experiments were repeated in the presence of soluble and insoluble carbon sources. M9 media were supplemented with 0.25% w/v casamino acids and varying concentrations of yeast extract, sodium acetate, glucose, sucrose (w/v), or glycerol (v/v): 0.00%, 0.05%, 0.25%, and 0.50%. Additionally, n-heptadecane (0.1% v/v) was tested under similar conditions. Flasks were inoculated with 1 mL of an overnight culture of *Gordonia* sp. iso11, with a minimum OD_600_ of 0.600, and incubated under aerobic conditions at 37 °C for 28 days, with the same media replenishing regime as described above.

### 2.6. First-order rate

Given that substrate utilization by microbes is typically fitted to a first-order rate model [32, 33], the rate constant (*k*) was used to compare experimental conditions:

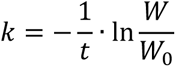

where *t* = time in days, *W* = mass of substrate after *t* time, and *W_0_* = mass of substrate at *t* = 0.

### 2.7. Quantification of biofilms formed by Gordonia sp. iso11

Biofilm formation was quantified in a microtiter-plate assay based on an absorbance method developed by Stepanović et al. [34]. Cells were grown in Nunc™ 96-well polypropylene plates (Thermo-Scientific, Rochester, NY). Sterile M9 media with varying carbon, iron, and surfactant contents were prepared and 190 μL were transferred into each well, with one column per condition. The wells at the extremities of the columns were not measured to ensure uniform testing conditions across the column [35] (i.e. six replicates of each condition). Each well was inoculated with 10 μL of pure bacterial culture suspended in the corresponding medium, to a total of 200 μL per well. Cells were grown aerobically at 37 °C for 18 h, after which the wells were emptied and refilled with 125 μL 0.1% w/v crystal violet solution and allowed to stain for 10 minutes. The wells were subsequently emptied and thoroughly rinsed by repeated immersion in deionised water and dried overnight. The crystal violet stain in each well was resolubilised in 125 μL 30% v/v acetic acid solution and absorbance readings were taken using a CLARIOstar Plus plate reader (BMG Labtech, Ortenberg, DE) at 570 nm (after 5 s double-orbital shaking at 500 rpm); absorbance readings were correlated to biofilm formation.

The ratio of planktonic cells to surface-attached cells was determined by OD_600_ measurements of the media in which the cells were cultivated in the 96-well plate. The 200 μL of media were transferred to a fresh 96-well plate prior to this reading to prevent interference by surface-attached cells. Ratios were calculated according to:

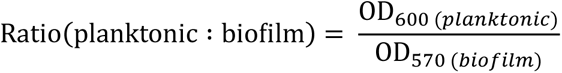

Furthermore, viable counts of plastic-attached cells were taken at the endpoint of the degradation experiments. Films were placed into 5 mL of carbon-free M9 medium in 50 mL conical tubes and sonicated at ambient temperature for 45 minutes in a Branson M Series Ultrasonic Cleaning Bath (Branson Ultrasonics, CT, USA). The resulting suspension was serially diluted, streaked onto M9 agar, grown aerobically for 24 h at 37 °C, and colonies counted. CFU cm^-2^ for a film with a total surface area of 16 cm^2^ was calculated according to:

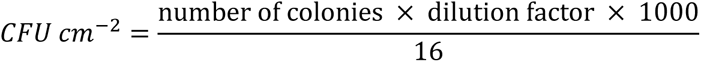

### 2.8. Determination of biosurfactant production

Microbial surfactant production was determined by measuring the contact angle of a 10 μL droplet of the supernatant of microbial growth medium on an untreated polypropylene surface. The static sessile drop method [36] was used on an Attension Theta Optical Tensiometer (Biolin Scientific, Gothenburg, SE). The supernatant was obtained by centrifugation of cell cultures for 3 minutes at × 12 000 g.

### 2.9. Whole genome sequence analysis

*Gordonia* sp. iso11 was grown in M9 broth, containing 0.5% w/v D-glucose and 0.25% w/v yeast extract, at 37 °C for 48 h. Genomic DNA (gDNA) extraction was carried out on *Gordonia* sp. iso11 using GenElute™ Bacterial Genomic DNA Extraction Kit (Merck KGaA, Darmstadt, DE), using the standard protocol for Gram-positive bacteria. Sequencing of purified gDNA was carried out by MicrobesNG (Birmingham, UK) using an Illumina sequencing platform with 30x coverage. Reads were trimmed using Trimmomatic, producing non-overlapping paired ends at the end of each contig [37]. Annotation was carried using PROKKA [38] and the NMPDR RAST (Rapid Annotation using Subsystem Technology) pipeline [39], and visualised using Proksee [40].

The NCBI BLAST server (tBLASTn) was used to find any significant alignments between the whole genome sequence and known protein sequences based on the published UniProt assessions as listed on the PlasticDB database (https://plasticdb.org/) [41].

### 2.10. AlkB protein expression

Plasmids (pET28a(+)) were synthesized with the *alkB* gene insert from *Gordonia* sp. iso11 at the BamHI and HindIII restriction sites, with the TGA stop codon at the HindIII restriction site. The recombinant plasmids were used to transform *Escherichia coli* BL21 (DE3) cells. Cells transformed with blank and recombinant plasmids were grown overnight in 1 L LB broth containing 50 mg mL^-1^ kanamycin in a 2 L Erlenmeyer flask, at 37 °C with 100 rpm shaking, until the OD_600_ reached 0.8, after which the cells were cooled to room temperature. Cells were induced with 100 μM IPTG and stirred for 18 h at room temperature. Following induction, the cultures were centrifuged at 6000 × g for 20 minutes. The pellets (from 200 mL aliquots) were resuspended in 25 mL phosphate-buffered saline (PBS) solution and lysed by sonication (frequency >20 kHz) in conical tubes using a probe sonicator, with cooling cycles (cycles of 20 s sonication, followed by 10 s cooling, for 3 minutes). The resulting suspensions were centrifuged (6000 × g for 10 minutes) to separate the soluble (supernatant) and insoluble (cell debris) fractions. Cell pellets were prepared into powder by freeze drying.

SDS-PAGE was carried out on both fractions to detect AlkB protein expression. 5 μL denaturing buffer (Laemmli) were added to 25 μL of each sample, along with 10-fold dilutions, (i.e., 5 μL Laemmli buffer, 2 μL of sample, and 18 μL water). These were incubated at 95 °C for 10 minutes, and the loaded into gel (Invitrogen™ NuPAGE™ 4-12% Bis-Tris 1.0 mm; Thermo-Fisher Scientific), along with the protein standard (Invitrogen™ SeeBlue™ Plus2; Thermo-Fisher Scientific). Electrophoresis was carried out at 200 V in NuPAGE MES/MOPS SDS-PAGE buffer (Thermo-Fisher Scientific) for 45 minutes. The gel was stained by immersion in Coomassie brilliant blue solution for 1 h, and then de-stained by immersion in an aqueous mixture of 40% (v/v) methanol and 10% (v/v) acetic acid for 2 h. The gel was imaged digitally immediately following de-staining.

## 3. Results and discussion

### 3.1. Screening and enrichment of plastic-colonizing microbes

The screening and enrichment methods yielded a total of 12 plastic-colonizing bacteria which displayed growth during the 28-day enrichment with various synthetic plastics as the sole carbon and energy source. Bacteria were identified by 16S rRNA gene sequencing and alignment using NCBI BLAST (Figure 1). *Gordonia* sp. iso11 and *Ralstonia* sp. iso23 were isolated from the peat bog wetland and were recovered from PP and PS, respectively (Figure 1). The remaining isolates, *Arthrobacter* sp. iso16, *Rhizobium* sp. iso34, *Enterobacter* sp. iso4, and several *Pseudomonas* spp. were isolated from domestic compost and were all recovered from PS following the 28-day enrichment (Figure 1).

**Figure 1.**
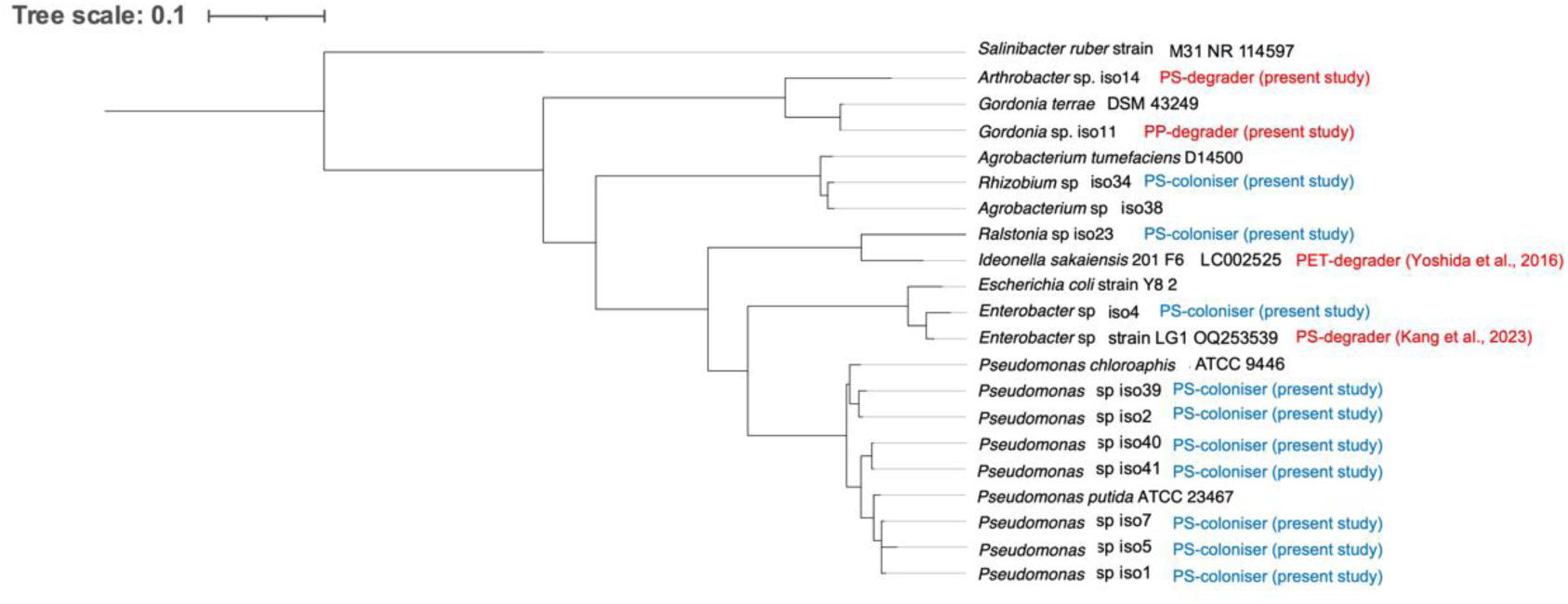
Maximum likelihood phylogenetic tree displaying the 16S rRNA gene distances between isolates calculated from 1000 bootstrap trials, based on nucleotide substitutions per sequence position. Other plastic-degrading microbes are included, along with closely related type strains; *Salinibacter ruber* (NR_114597) used as an outgroup root.

Whilst the growth conditions and experimental design may have facilitated fungal and/or archaeal growth, morphology and 16S sequencing provides no evidence of any proliferation of these microbes. It is noted that the presence of some species may have been inhibitory to the proliferation of others in the early phases of the screening experiment. Therefore, some loss in the microbial diversity and/or flux in the microbial community composition is to be expected towards the end of the 28-day incubation [42]. Regardless, the 28-day incubation period is necessary to ensure selection of plastic-colonizing microbes in a nutrient-deplete environment, considering the recalcitrance of polymeric substrates (owing to their high degree of hydrophobicity [18], and expected slow rates of degradation [19]).

*Arthrobacter* sp. iso16 displayed 98.80% identity with *Arthrobacter* sp. strain TS-15 (MK459547) [43]. *Arthrobacter* spp. are Gram-positive bacteria, common in soil [44]. In previous studies, *Arthrobacter* spp. are isolated on hydrophobic substrates including dibenzofuran, with evidence of degradation of related hydrophobic compounds dibenzo-p-dioxin and 2-monochlorodibenzo-p-dioxin [45], both of which are emerging environmental pollutants.

*Gordonia* sp. iso11 is one of two isolates from the nutritionally complex peat soil sample to colonize polypropylene where it was the sole carbon source (Figure 1); it displayed 99.72% 16S gene sequence identity with *Gordonia terrae* strain 3612 (CP096585). Species from the genus *Gordonia* have been regarded as ‘metabolically diverse’ due to their ability to degrade and assimilate a wide variety of carbon sources [25]. *Gordonia* species are prevalent in diverse habitats, including mangrove and terrestrial ecosystems [28]. A polystyrene degradation study on species closely related to *Gordonia sihwensis* found PS degradation rates of up to 7.7% in a 28-day period [46]. Other isolation studies have characterized *Gordonia* spp. degrading PET [47] and PE film [48]. *G. terrae* has been isolated from oil-polluted soils, forming biofilms on long chain alkanes [49]. This is likely attributed to the presence of *alkB* and cytochrome P450 genes [50], and/or other monooxygenase-encoding genes which are common in *Gordonia* species [27, 51, 52].

In general, based on the scientific literature, the isolation study appears to have yielded hydrocarbon-degrading genera that can be isolated from diverse niches.

### 3.2. Mass loss of PS and PP films by Arthrobacter sp. iso16 and Gordonia sp. iso11

Microbial degradation of high-purity plastic films, free from any additives, was measured by mass loss, where they were the sole carbon and energy source available. The technique has been widely employed in other studies throughout the literature [20, 53]. Each isolate was assayed for degradation activity on the plastic from which they were enriched and isolated. Of these isolates, *Gordonia* sp. iso11 and *Arthrobacter* sp. iso16 displayed the highest rates of degradation across the triplicate experiments. *Gordonia* sp. iso11 degraded PP film by up to 22.8% (corresponding to 6.52 mg); *Arthrobacter* sp. iso16 degraded PS film by up to 19.5% (4.67 mg; Figure 2).

**Figure 2.**
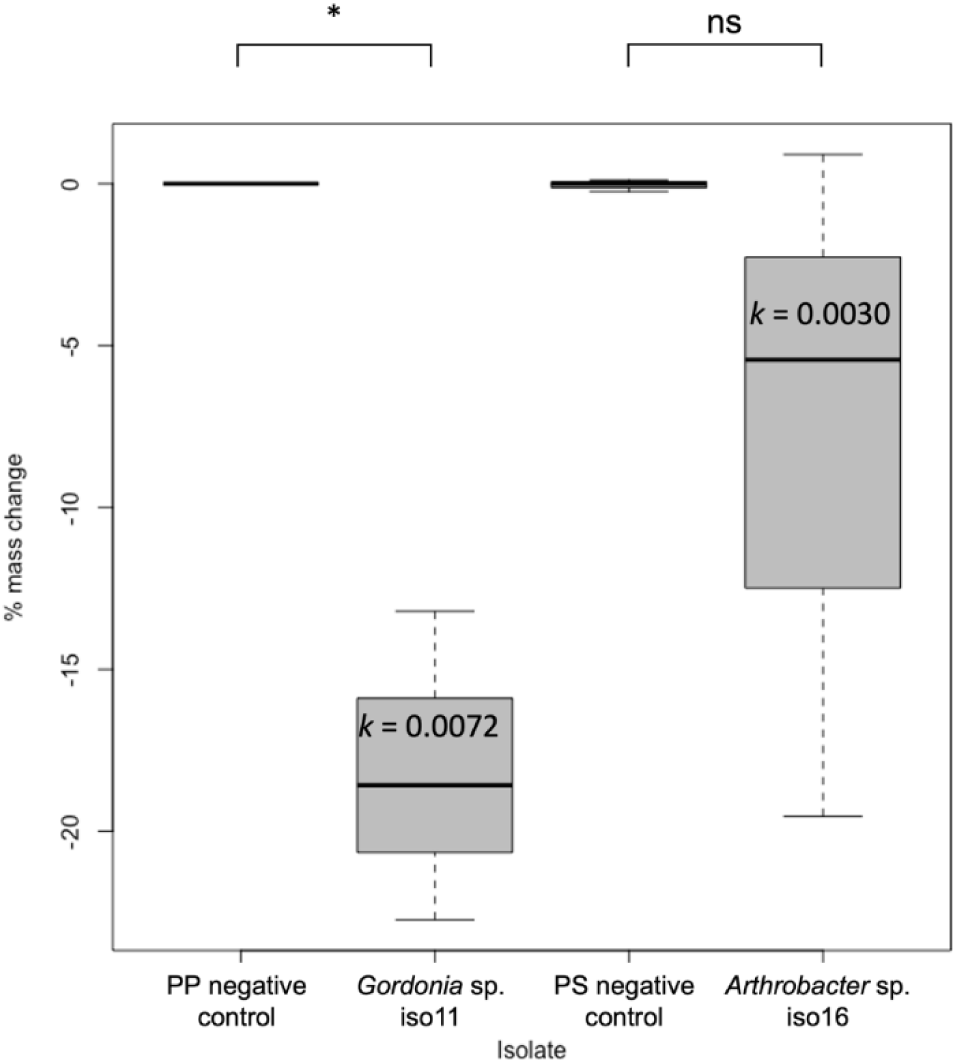
Percentage mass change of plastic films following 28-day incubation with *Gordonia* sp. iso11 (with PP), and *Arthrobacter* sp. iso16 (with PS) in M9 medium, supplemented with 0.25% w/v casamino acids. Plotted values are the median of triplicate measurements, whiskers represent minimum and maximum values, and the bottom and top of the box show the first and third quartiles, respectively. First-order rate constants (*k*) were calculated from mean data (n = 3). Asterisks (*) indicate significance; one-way ANOVA, followed by Tukey HSD post-hoc test, *p* < 0.05.

Use of one-way ANOVA demonstrated that PP degradation by *Gordonia* sp. iso11 is significant when compared with the control (p < 0.05), whereas the high variability in PS degradation by *Arthrobacter* sp. iso16 led to statistical insignificance (Figure 2).

*Gordonia* sp. iso11 displayed the highest rates of PP degradation, with rates of 22.73% over a 28-day period (0.23 ± 0.02 mg d^-1^; *k* = 0.0072) (Figure 2); the highest degree of degradation observed in this study. In terms of rate, this is comparable to other studies involving soil bacteria. In *Enterobacter hormeachi* WX-2, aerobic PP degradation rates of 4.7 ± 0.1% were observed over a 60 d period [20], and in *Brevibacillus agri* btDSCE02, 27% ± 2% mass loss was observed following 140 day aerobic incubation with at 50 °C [54].

*Arthrobacter* sp. iso16 displayed a comparable degree of PS degradation, up to 19.54% (0.17 ± 0.02 mg d^-1^; *k* = 0.0030). Other mass loss studies on high-purity PS have reported rates of 1.52% by *Acinetobacter johnsonii* JNU01 over 30 days [55], 7.4% over 60 days by *Exiguobacterium* sp. YT2, isolated from mealworms [56], and up to 12.4% by an *Enterobacter* sp. with decabromodiphenyl oxide and antimony trioxide additives [57]. Kim et al. [55] attributed PS degradation by *A. johnsonii* JNU01 to a 10-fold increase in alkane-1-monooxygenase (*alkB*) gene expression.

### 3.3. Degradation of PP by Gordonia sp. iso11 in the presence of other carbon substrates

Addition of other carbon substrates to the microcosm experiments reflects better on the *in-situ* conditions; it provides insights to biodegradation performance where substrates are present which are available to the microbe in nutritionally complex niches. A range of carbon sources with varying bioavailability were tested. These were sodium acetate, which may be assimilated in some extremophilic microbes [58], yeast extract, glucose, sucrose (w/v), or glycerol (v/v) at concentrations of either 0.05% or 0.5%.

Where there was an addition of soluble carbon (Figure 3), yeast extract and glycerol were the only additional carbon sources where PP degradation occurred. Yeast extract was chosen because it contains highly diverse carbon sources; glycerol is a compatible solute which, in bacteria, mitigates oxidative stresses [59], such as those involved by the release of free radicals in plastic breakdown. Like the components of yeast extract, glycerol can be metabolized and mineralized by the cell to support proliferation, to a limited extent (μ = 0.01 h^-1^), in some *Gordonia* spp. [60].

**Figure 3.**
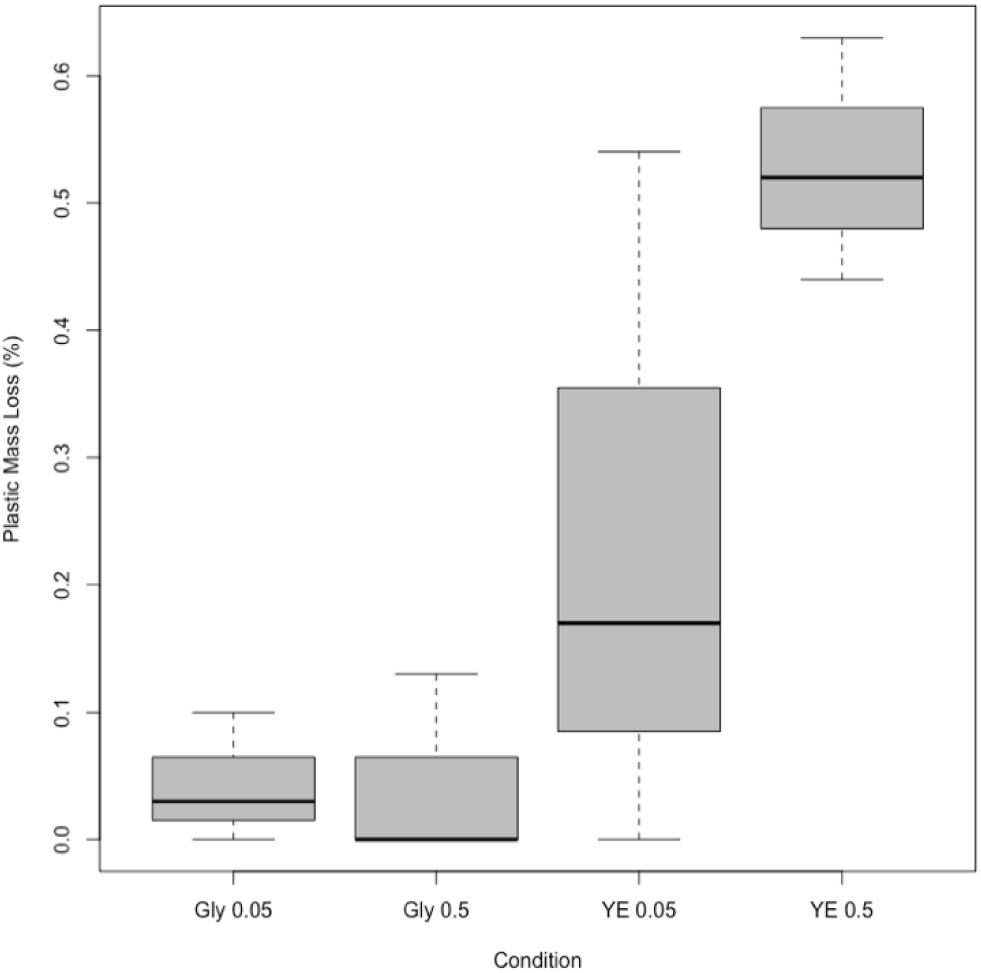
Percentage mass loss of PP by *Gordonia* sp. iso11 with various soluble carbon sources. <u>Abbreviations</u>: Gly 0.05 – glycerol 0.05% v/v; Gly 0.5 – glycerol 0.5% v/v; YE 0.05 – yeast extract 0.05% w/v; YE 0.5 – yeast extract 0.5% w/v.

Whilst PP degradation was observed under these conditions, it was significantly reduced when compared to the experiments with carbon-free M9 medium; rates of up to 0.63% over 28 d was observed in M9 supplemented with 0.5% w/v yeast extract (Figure 3), a significant reduction when compared to the 22.73% observed in carbon free M9 (Figure 2). A reduction in yeast extract (to 0.05% w/v), resulted in a decrease in degradation rates, and markedly increased the variability of the data compared to the 0.5% w/v yeast extract (Figure 3). It is likely that *Gordonia* sp. iso11 cells preferentially metabolize the components of yeast extract, and this metabolic support does not in any way upregulate genes for enzyme systems involved in plastic degradation. Virtually no degradation activity was observed in the incubations with glycerol (Figure 3).

Given that soluble carbon sources do not cause any increased rates of plastic degradation, and indeed is inhibitory to it, further experiments were carried out to determine whether the hydrocarbon-degrading apparatus in *Gordonia* sp. iso11 could be differentially expressed by environmental changes. Plastic mass loss experiments were conducted in M9 medium (supplemented with 0.25% w/v casamino acids and 0.003% w/v iron(III) chloride) containing 0.1% v/v n-heptadecane (C_17_), with or without the addition of 0.2% v/v Tween-80 non-ionic surfactant.

The data indicate mass loss of PP of 12.34% - 15.67% by *Gordonia* sp. iso11 with n-heptadecane for 28 d (Figure 4a). Maximum rates of degradation were increased in the presence of Tween-80 up to 22.63%, however this was coupled with a much higher degree of variability (7.56% - 22.63% mass loss). The results of PP mass loss with and without surfactant and n-heptadecane are significant compared to the negative control (*p* < 0.05) (Figure 4a). However, no significant change to the rate of degradation of PP compared to incubation in carbon-free M9 medium (Figure 4a). Incubation of *Gordonia* sp. iso11 with n-heptadecane resulted in a ∼6-fold increase in plastic-attached cells, when compared to the incubations in carbon-free M9 medium and in M9 medium containing Tween-80 (Figure 4b). There was no noteworthy change in plastic-attached cell counts between the incubations with n-heptadecane and Tween-80, and carbon-free M9 medium. The viable counts of the planktonic cells (Figure 4c) throughout the 28-day incubation followed similar trends, regardless of media condition: a peak cell count upon initial dense inoculation with *Gordonia* sp. iso11 (2-5×10^11^ CFU mL^-1^), a slight decrease at day 7 (2×10^9^ - 1×10^11^ CFU mL^-1^), a marked decrease at day 14 (2×10^7^ - 6×10^8^ CFU mL^-1^), an increase at day 21 (2×10^7^ - 2×10^10^ CFU mL^-1^), followed by a highly variable decrease at the endpoint (4×10^5^ - 2×10^8^ CFU mL^-1^). The variability in these data (in terms of standard deviation from the mean) generally increased as the experiment progressed (Figure 4c).

**Figure 4.**
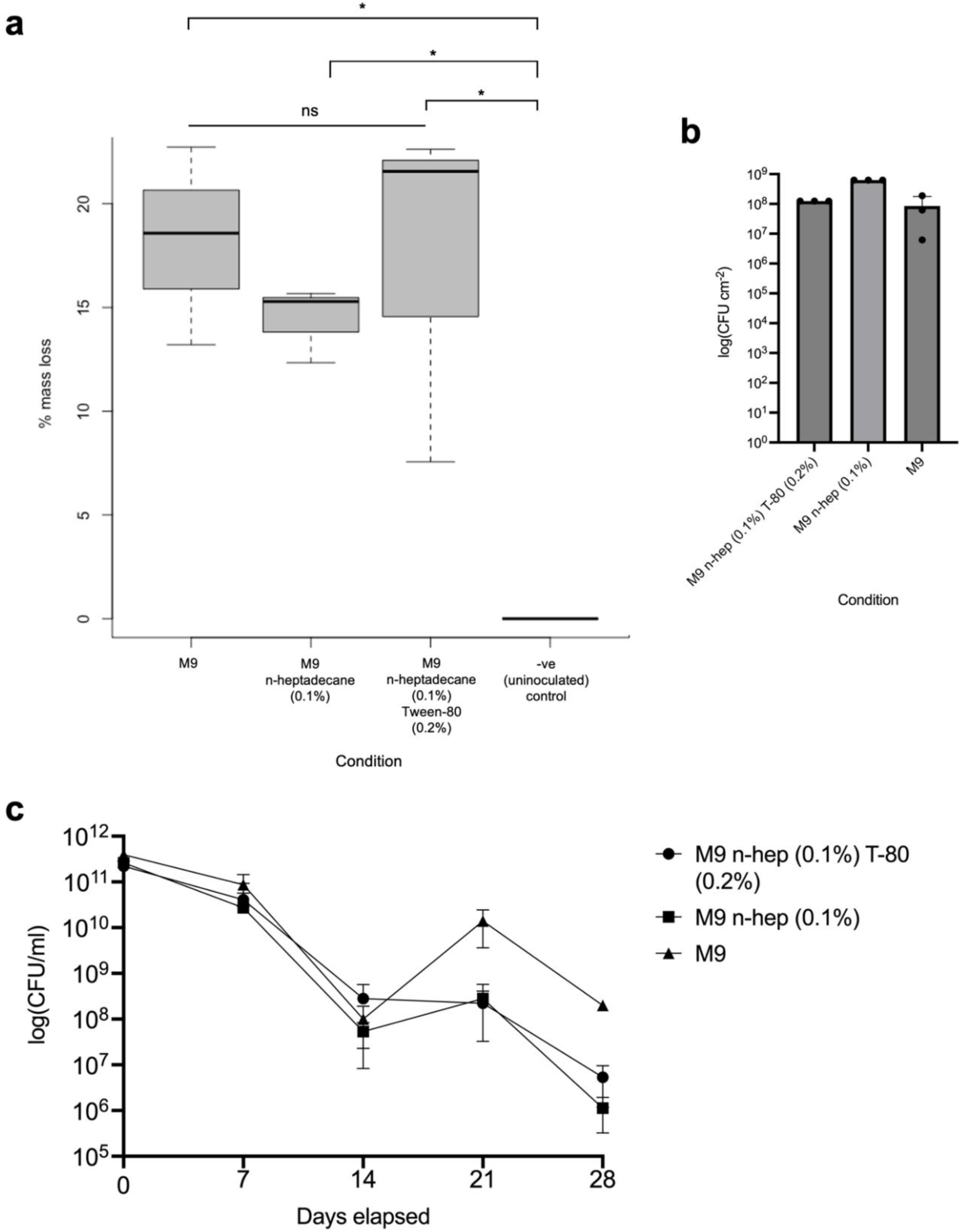
Degradation of PP by *Gordonia* sp. iso11 in the presence of n-heptadecane. **a** percentage mass loss of PP film with and without Tween-80 surfactant; **b** total viable counts of plastic-attached microbes after 28 days incubation (CFU cm^-2^); **c** viable counts of planktonic cells throughout the 28-day incubation. Asterisks indicate significance: one way ANOVA, followed by Tukey HSD test, *p* < 0.05. <u>Abbreviations:</u> n-hep – n-heptadecane; T-80: Tween-80. All media contain 0.25% w/v casamino acids.

Overall, incubation with n-heptadecane greatly decreased the variability of the data but did not significantly change the rate of degradation (Figure 4a). The rationale for these experiments is that the presence of an alkane may upregulate hydrocarbon-degrading enzymes in *Gordonia* sp. iso11. In a study on *Pseudomonas aeruginosa* PAO1 and RR, increased expression of an alkane hydrolase system (involving genes *alkB1, alkB2, alkG1, alkG2, alkT*) was observed in the presence of C_10_ to C_22_/C_24_ alkanes [61]. Another study used hexadecane to enrich a consortium predominately composed of *Gordonia* spp., yielding PE-degrading *Gordonia polyisoprenivorans* B251 [62]. Therefore, it was hypothesized that a similar upregulation may occur in the hydrocarbon-degrading genes in *Gordonia* sp. iso11 when it is incubated with a hydrocarbon in liquid phase. Upregulation of these genes may enhance plastic degradation if the proteins involved with plastic breakdown are more highly expressed and available to approach the surface of the polymer. For this purpose, n-hexadecane was chosen for two key reasons: (i) its low volatility at the incubation temperature of 37 °C, and (ii) that its molecular mass falls within the range of peak alkane hydrolase expression of all long-chain hydrocarbons tested in the Marin et al. study [61]. Alkanes and polyolefins share characteristics pertinent to their low degradation rates by microbes (low bioavailability and high hydrophobicity). They have a backbone composed of relatively strong C-C covalent sigma bonds, which require large energy input to break (∼350 kJ mol^-1^ at 298 K [63]). Despite this, alkane degradation by bacteria and specifically several *Gordonia* spp. is well characterized [50, 52, 64]. Given the previously observed plastic degradation activity, it is possible that *Gordonia* sp. iso11 possesses a similar molecular apparatus. Addition of Tween-80 was carried out to determine whether it enhanced plastic degradation by decreasing the interfacial tension between the aqueous phase and solid plastic, which may facilitate the initial interaction of microbial enzyme systems and polymer surface.

The results obtained from the Tween-80 experiment did not provide evidence of increased degradation activity in comparison to incubation with only PP film. However, when compared to mass loss rates of the different soluble carbon sources tested (Figure 3), these data demonstrate a marked increase in rates, which may be attributed to the cells’ resources being directed towards mechanisms capable of degrading hydrocarbons. If any upregulation of genes responsible for hydrocarbon degradation occurred, it would have facilitated mineralization of the alkane as well as the plastic, and therefore a more-optimal rate of plastic mineralization could not have occurred as the metabolic apparatus were mineralizing two substrates simultaneously.

The marked increase in plastic-attached cells observed in the incubations with n-heptadecane is attributed to a dipartite action: (i) an interaction between the two non-polar phases (i.e., the liquid alkane and the solid plastic), on which cells are interacting, and (ii) changes to the protein expression at cell surfaces changes the cell surface hydrophobicity and increases hydrophobic interactions between hydrophobic residues on the cell surface and substrate [65]. This increase is not observed in the incubation with Tween-80, which may be expected given that the 0.2% v/v concentration used exceeds that of that observed to inhibit biofilm formation (0.1% v/v) [66]. However, use of surfactant did not cause a decrease in the microbial attachment when compared to the carbon-free M9 (Figure 4b). It is likely that cellular adaptation to the hydrophobic environment has a critical influence on these results. As mentioned above, changes in the protein expression at cell surfaces make them more (or less) hydrophobic. If use of a surfactant decreased the hydrophobicity of the alkane and/or plastic surface, it may have reduced the cellular response to the presence of the hydrophobic substrates, reducing the cell surface hydrophobicity, which correlates directly with attachment [67].

### 3.4. Effect of nutrient conditions on biofilm formation and growth rate of Gordonia sp. iso11

Results from the biofilm assay indicate that there are trends in biofilm formation across media conditions (Figure 5). All readings were blanked against an uninoculated column of wells in a 96-well plate containing M9 medium to remove the effect of background staining on the bottom and sides of the well. Based on means of six replicates, the highest biofilm formation was observed in 0.25% w/v sodium acetate with a mean absorbance of 0.104 (SD = 0.010) at 550 nm. The weakest biofilm formation was observed following incubation with 0.05% w/v yeast extract in M9 medium, with a mean absorbance of 0.029 (SD = 0.014). M9 medium with no additives was also inoculated with *Gordonia* sp. iso11 and used as a point of statistical comparison. Whilst no condition significantly enhanced biofilm formation, one-way ANOVA, followed by Dunnett’s multiple comparisons test indicated that 0.05% w/v yeast extract and presence of 0.003% w/v Fe^3+^ significantly inhibited biofilm formation (p < 0.05) (Figure 5). This is particularity pertinent to hydrocarbon degradation, given that Fe^3+^ is a common enzyme cofactor which binds four histidine residues in AlkB proteins [12, 68]. In the case of *Gordonia* sp. iso11, excess of Fe^3+^ could result in lower rates of attachment to plastics and ultimately lower their rates of degradation.

**Figure 5.**
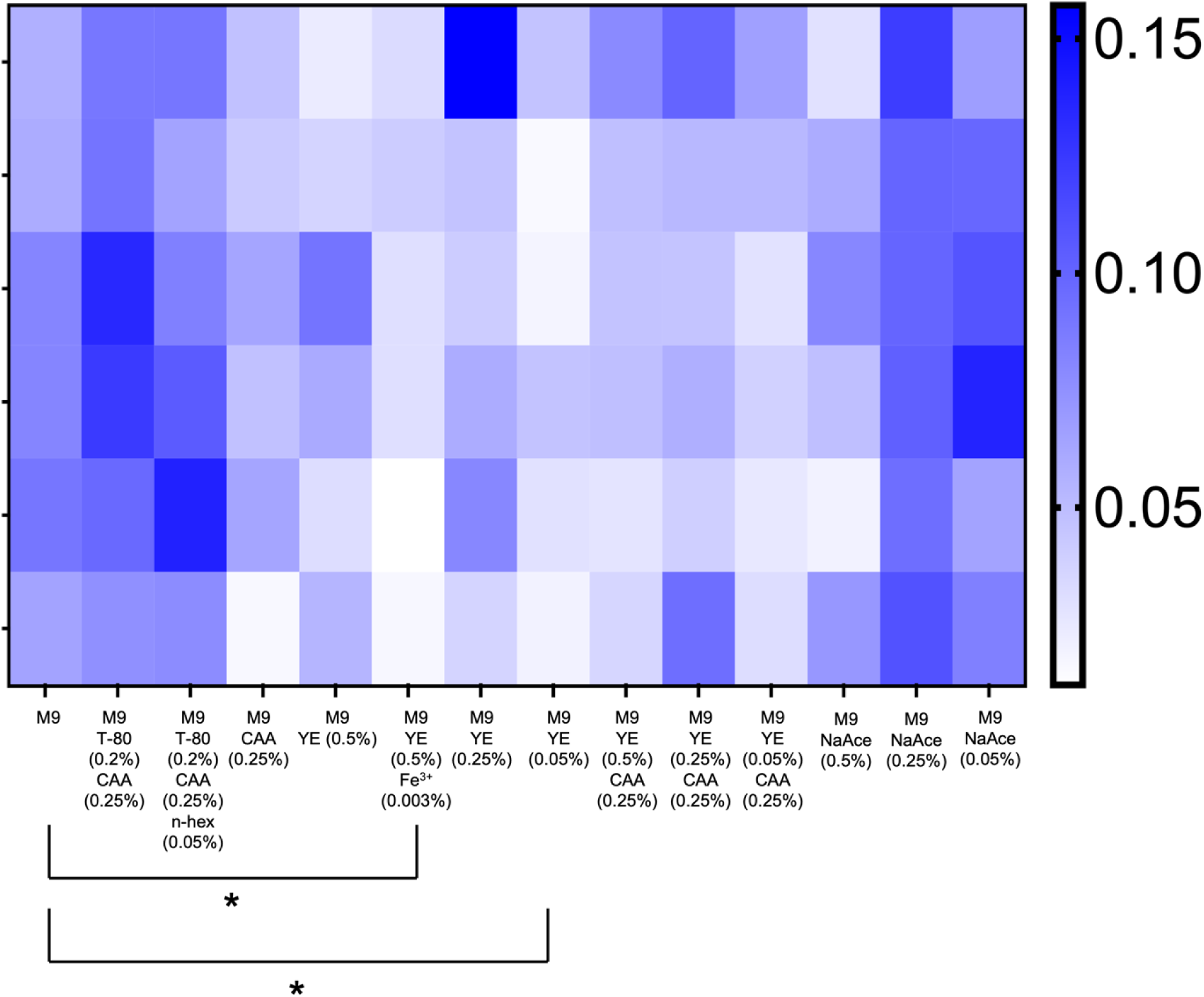
Absorbance values (570 nm) of resolubilized crystal violet, indicating trends in biofilm formation by *Gordonia* sp. iso11 after 18 h aerobic growth under varying media conditions. Asterisks indicate pairwise significance: one-way ANOVA, followed by Dunnett’s multiple comparisons test, *p* < 0.05. <u>Abbreviations</u>: T-80 – Tween-80; CAA – casamino acids; n-hex – n-hexadecane; YE – yeast extract; NaAce – sodium acetate. Percentages are w/v, except T-80 and n-hex, v/v.

Interestingly, provision of 0.05% w/v yeast extract also caused significant inhibition of biofilms (Figure 5), especially when compared to the completely nutrient-deplete M9. In general, incubation with yeast extract at any concentration resulted in poorer biofilm formation, with the use of 0.05%, 0.25%, and 0.50% w/v yeast extract resulting in comparable mean absorbance values of 0.029 (SD = 0.014), 0.070 (SD = 0.046), and 0.049 (SD = 0.025), respectively. Some microbes exhibit increased biofilm formation as part of a transcriptional response to conditions causing nutrient stress [69]. Provision of yeast extract to the culture medium adds many complex bioavailable components such as organic carbon, amino acids, peptides, phosphates, fatty acids, and metal ions [70]. Therefore, the marked reduction in biofilm formation with yeast extract can be attributed to a multi-faceted effect: the presence of trace metal ions which may have disrupted biofilms, and that the cells were not sufficiently nutrient-stressed to cause the formation of a dense biofilm, compared to the other conditions tested. By this reasoning, it can be inferred that the other conditions introduce a nutrient stress to this bacterium.

Ratios were calculated for each condition to demonstrate the proportion of cells which were planktonic in the media, versus those that were attached to the bottom and sides of the wells via biofilm formation. The highest proportion of planktonic cells, compared to those in a biofilm, was observed in 0.5% w/v yeast extract with 0.003% w/v Fe^3+^, with a planktonic cell population up to 18.8 times higher than those in a biofilm (mean = 9.73; SD = 5.19) (Figure 6a). This correlates well with the data indicating significantly less biofilm production (Figure 5). Whilst the greatest proportion of cells cultivated with Fe^3+^ are planktonic, growth rates slowed from 0.14 h^-1^ to 0.09 h^-1^ in the presence of Fe^3+^ (Figure 6b). This demonstrates that under these more nutrient-rich conditions, addition of Fe^3+^ almost completely inhibits the production of biofilms. Addition of 0.25% w/v casamino acids to varying concentrations of yeast extract greatly reduced the variation in the ratios, with means of 3.55 (SD = 1.27), 3.21 (SD = 1.41), and 3.52 (SD = 1.21) for yeast extract concentrations of 0.05%, 0.25%, and 0.50% w/v, respectively (Figure 6a). This treatment also slightly reduced mean growth rates (Figure 6b).

**Figure 6.**
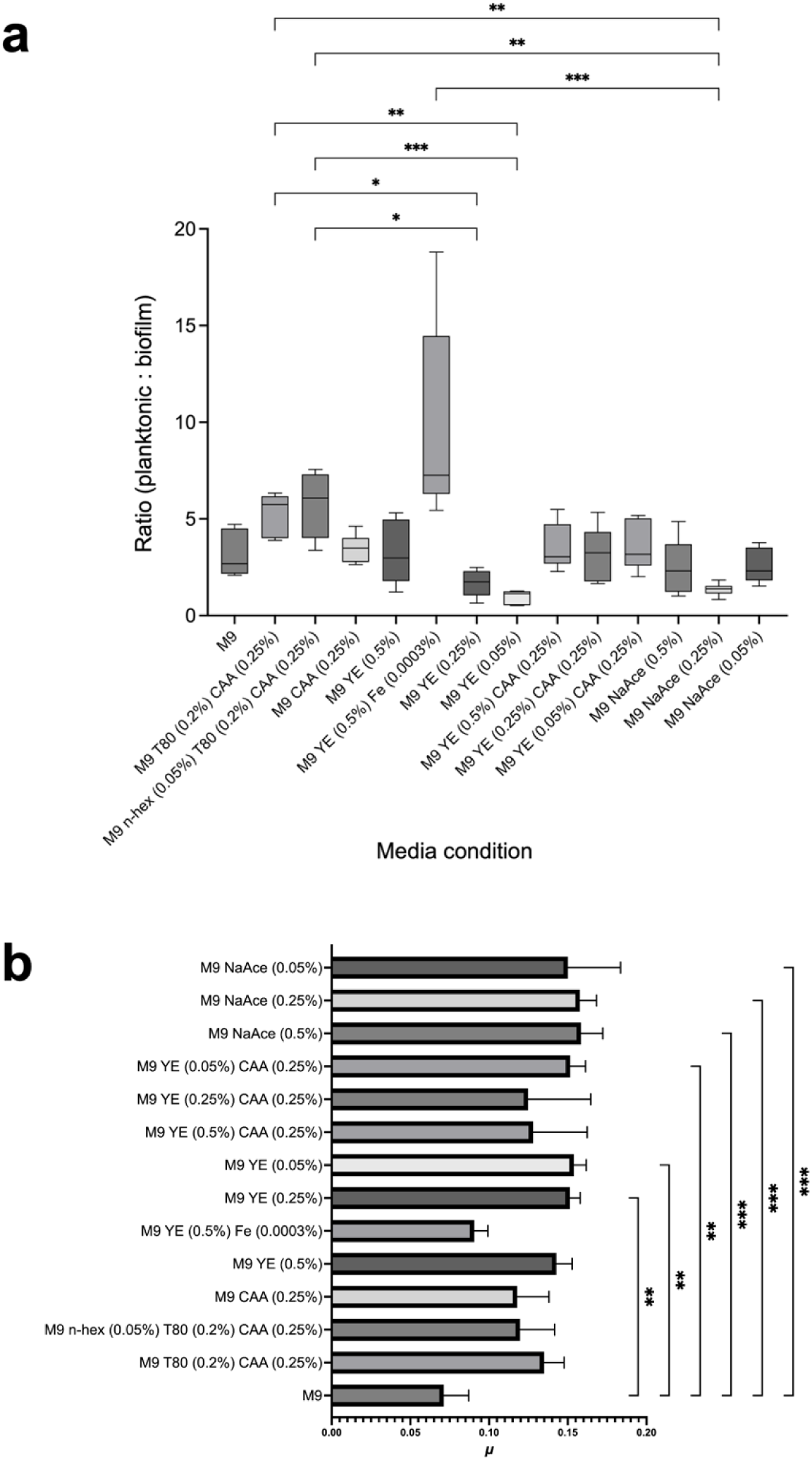
a. Ratios of planktonic *Gordonia* sp. iso11 cells to biofilm-attached cells after 18 h aerobic growth under varying media conditions. Asterisks indicate all pairwise significance according to Dunn’s multiple comparisons test; **b** corresponding specific growth rates (*μ,* h^-1^), with pairwise comparisons to the M9 control; * *p* < 0.05, ** *p* < 0.005, *** *p* < 0.0005. <u>Abbreviations</u>: T-80 – Tween-80; CAA – casamino acids; n-hex – n-hexadecane; YE – yeast extract; NaAce – sodium acetate. Percentages are w/v, except T-80 and n-hex, v/v.

A considerable amount of growth occurred in the carbon-free M9 medium, with a mean of 0.07 h^-1^ (Figure 6b), despite there being no soluble carbon or nitrogen sources. A similar effect was observed in a study which investigated the effects of casamino acids and other nutrients on cell motility and surface adhesion, where an inoculated control composed of M9 medium displayed ∼100-fold increases in viable cell counts over a four day period [71]. Such growth in nutrient-deplete media is caused by metabolism of cell debris in the medium, and the transfer of small quantities of other carbon sources originating from the culture used as an inoculum [71].

*Gordonia* sp. iso11 displayed no growth in any concentration of glycerol. Similar results were observed from experiments on *G. polyisoprenivorans* VH2, which displayed a growth rate on 0.2% w/v glycerol of only 0.01 h^-1^ [60].

Whilst some studies on the degradation of polymeric substrates by bacteria have utilized a ‘clear zone’ methodology for observing the degradation of the on agar [72], members of the genera *Gordonia* and *Nocardia* do not exhibit the same behavior. Typically, they grow adhesively on the polymer surface in a more effective polymer-degrading mechanism [60, 72]. From this point of view, elucidating the biofilm-forming conditions by plastic-degrading microbes such as *Gordonia* sp. iso11 is critical to understanding how to improve its rates of polymer degradation. Since the microbe does not form clear zones on agar, it is unlikely that it secretes enzymes to degrade the polymer, as it would in a saprotrophic mechanism, but rather utilizes membrane-bound and/or transmembrane proteins requiring direct attachment to the substrate.

### 3.5. Biosurfactant production by Gordonia sp. iso11

Surfactants reduce the surface tension of liquids and will result in a reduction of the contact angle of liquid on a hydrophobic surface. Therefore, contact angles were measured and displayed as means across biological triplicate experiments (Figure S1).

A clear trend was observed in the contact angles of the supernatants following incubations with *Gordonia* sp. iso11, when compared to the blanks (Figure 7c). Growth in M9 with 0.5% w/v D-glucose for 18 h showed a marked reduction in the mean contact angle from 102.24° for the sterile M9 medium to 72.41° following 18 h incubation with *Gordonia* sp. iso11 (Figure 7c). There was very little change in contact angle observed when *Gordonia* sp. iso11 was incubated with PP or n-heptadecane (72.24° and 72.62°, respectively). A marked decrease in contact angle was observed when *Gordonia* sp. iso11 was incubated with Tween-80, with a mean of 57.37°, comparable to the mean of 54.33° observed with 0.2% v/v Tween-80 (Figure 7c).

**Figure 7.**
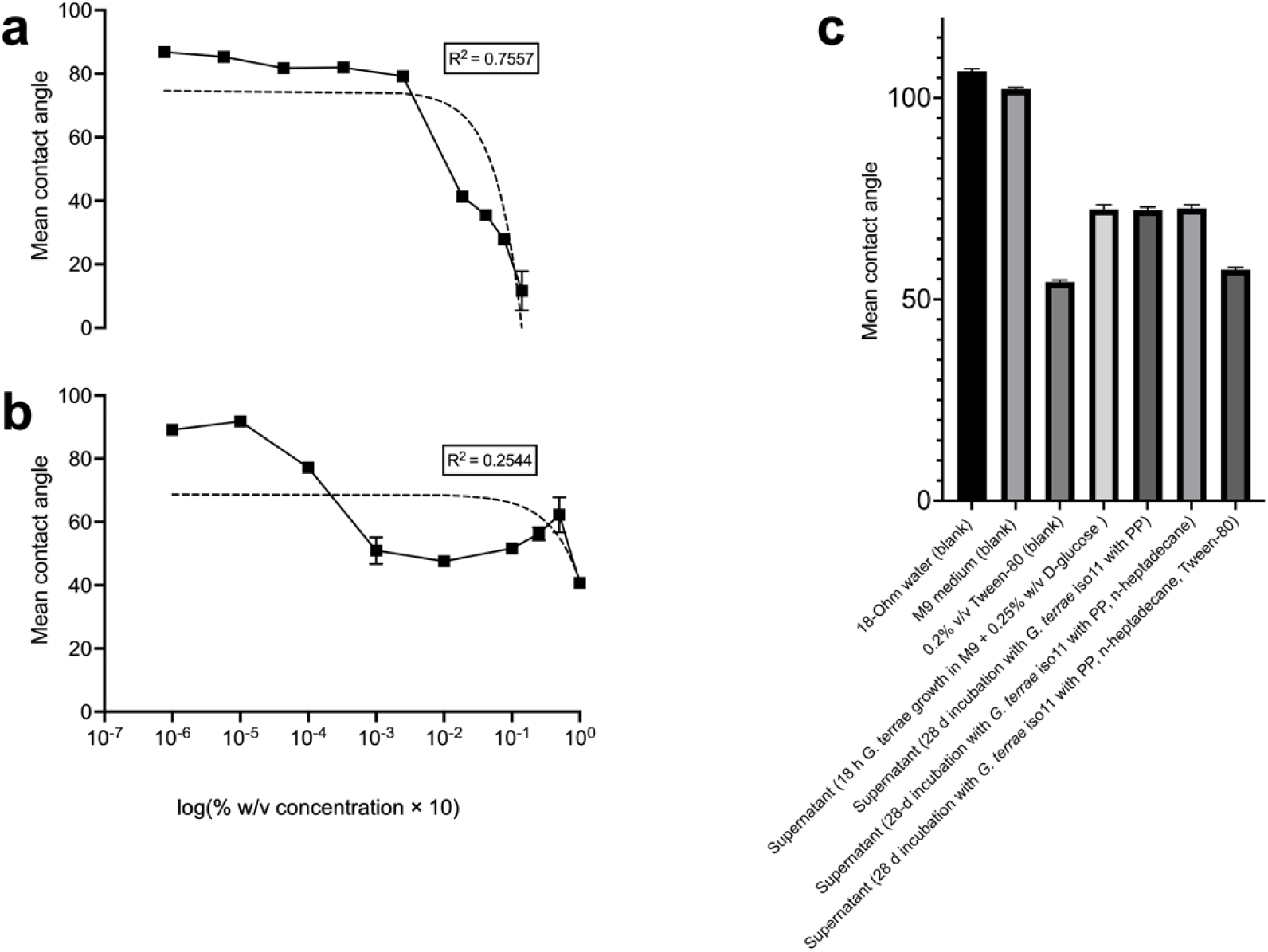
Measured contact angles of supernatants of degradation experiments with *Gordonia* sp. iso11, and surfactants of known concentration, on an untreated PP film. Simple linear regression of surfactant concentrations (**a** SDS, **b** Tween-80) and mean contact angles. R^2^ (goodness of fit) values displayed. **c** contact angles of *Gordonia* sp. iso11 PP degradation experiments, and blanks, on an untreated PP film. Error bars represent standard deviation from the mean.

The reduction in the mean contact angle of all media supernatants following incubations with *Gordonia* sp. iso11 indicates that the bacterium secretes surfactant-like compounds. These data also indicate that exposure to hydrophobic substrates causes no change to the amount of biosurfactant produced. Based on mean contact angles, the biosurfactants produced have a surface action equivalent of ∼2.89 g l^-1^ SDS (Figure 7a).

Biosurfactants are used by microbes to solubilize organic compounds, and increase their bioavailability [73]. Based on previous studies on *Gordonia* spp., such compounds may include fatty acids, bound to saccharides, with very high molecular weight (>10,000 Da) [74, 75]. It has been demonstrated that the high molecular weight fraction of compounds produced by *Gordonia amarae* SC1 are capable of emulsifying n-hexadecane [74]. Conversely, in similar conditions involving exposure to n-hexadecane, there was no biosurfactant production by two *Gordonia* strains: *G. terrae* S5 and *G. amicalis* S2 [49]. It is therefore likely that the production of biosurfactants is a strain-specific behavior.

The principle that surfactant concentration is proportional to the surface tension of a droplet was tested in this study using known concentrations of SDS and Tween-80 (with 10-fold dilutions) and fit to a linear regression model. The highest R^2^ (goodness of fit) value obtained was 0.7557 with SDS (Figure 7a), whereas a poor fit to this model was obtained with the same concentrations of Tween-80 (R^2^ = 0.2544) (Figure 7b). Given the variability of Tween-80 on the PP film, it could not be used for comparisons with the biosurfactant. By comparison, linear regression models were fit to similar data with both SDS and Tween-80, which gave R^2^ values of 0.9701 and 0.9639, receptively [76]. These were used to calculate critical micelle concentration (CMC) values of 1.84 g l^-1^ and 6.37 g l^-1^ for SDS and Tween-80, respectively [76]. From the data in this study, a CMC for SDS is 1.97 g l^-1^ (based on the point at which the concentration no longer causes a change in contact angle, according to the linear regression model).

### 3.6. Genome sequence analysis

*Gordonia* sp. iso11 harbors multiple genes coding for proteins capable of cleaving and/or oxidizing C-C and C-H bonds, including alkane-1-monooxygenase (AlkB) (Table S1) and cytochrome P450, and other enzyme systems which are involved in the transfer of electrons which enable these to function, including rubredoxin (RubA) (Figure 8). Several NADH-quinone reductases and alkyl hydroperoxide reductases are also present, which catalyze reactions involving hydrophobic substrates including quinone and alkyl groups (Figure 8).

**Figure 8.**
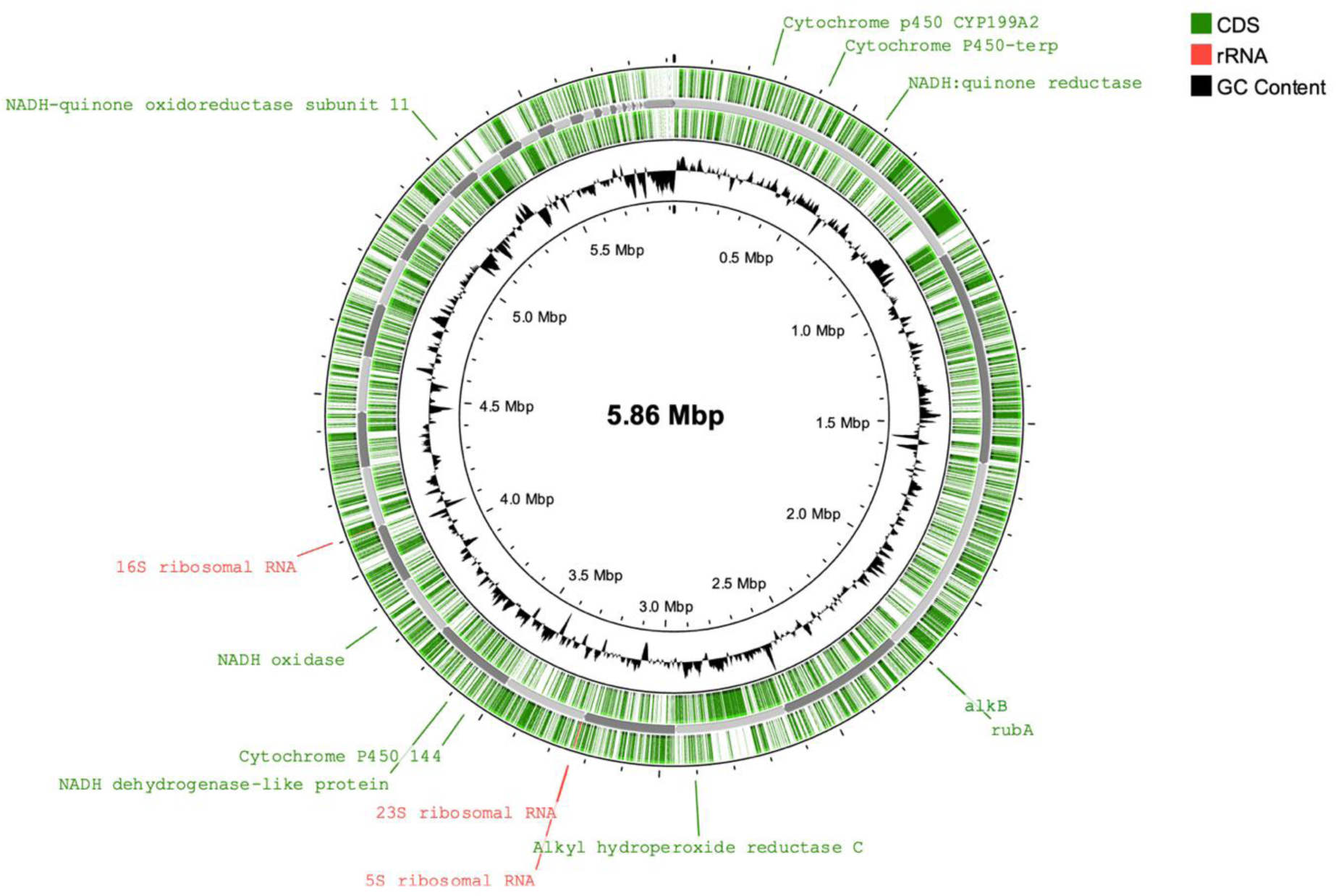
Genome map displaying the assembly and annotation of the sequencing data for *Gordonia* sp. iso11. The inner ring displays regions of high GC content. The green outer rings display the annotated ORFs according to PROKKA. The grey-shaded ring shows the contigs arranged by size. rRNA genes, and other genes pertinent to C-C cleavage, oxidation, and/or plastic degradation and functioning of AlkB, are labelled.

BLAST searches were carried out on the complete genome to determine the presence of genes coding for proteins homologous to those known to degrade synthetic plastics. Lists of percentage sequence identities are presented in Supplementary Table 1. Significant sequence identity was found in most of the sequences searched, with the highest being 75.06% for an alkane-1-monooxygenase from *Rhodococcus ruber* C208 (Table 1). In general, the highest sequence homology was found in genes coding for alkane-1-monooxygenase, for which there is empirical evidence demonstrating up 28.6% carbon mineralization of PE in 80 days [77].

### 3.7 Expression of Gordonia sp. iso11 AlkB

Based on annotations, *alkB* from *Gordonia* sp. iso11 has a size of 1236 bp, which will produce a protein of 46.88 kDa. Expression was carried out using the T7 expression system in *Escherichia coli* BL21 (DE3) cells, which were transformed with a synthesized AlkB/pET28a(+). The same experimental setup was carried out on a blank pET28a(+) plasmid. Whilst the presence of a single protein band, between 38 and 49 kDa, in the positive lanes (and its absence in the negative pET28a(+) lanes) of the polyacrylamide gel following SDS-PAGE (Figure 9) provides some evidence of successful protein expression under the induction conditions, the band is weak and indictive of low levels of expression. Further work must be carried out to enhance yields of the insoluble protein in order to obtain the milligram-quantities required for further isolation studies.

**Figure 9.**
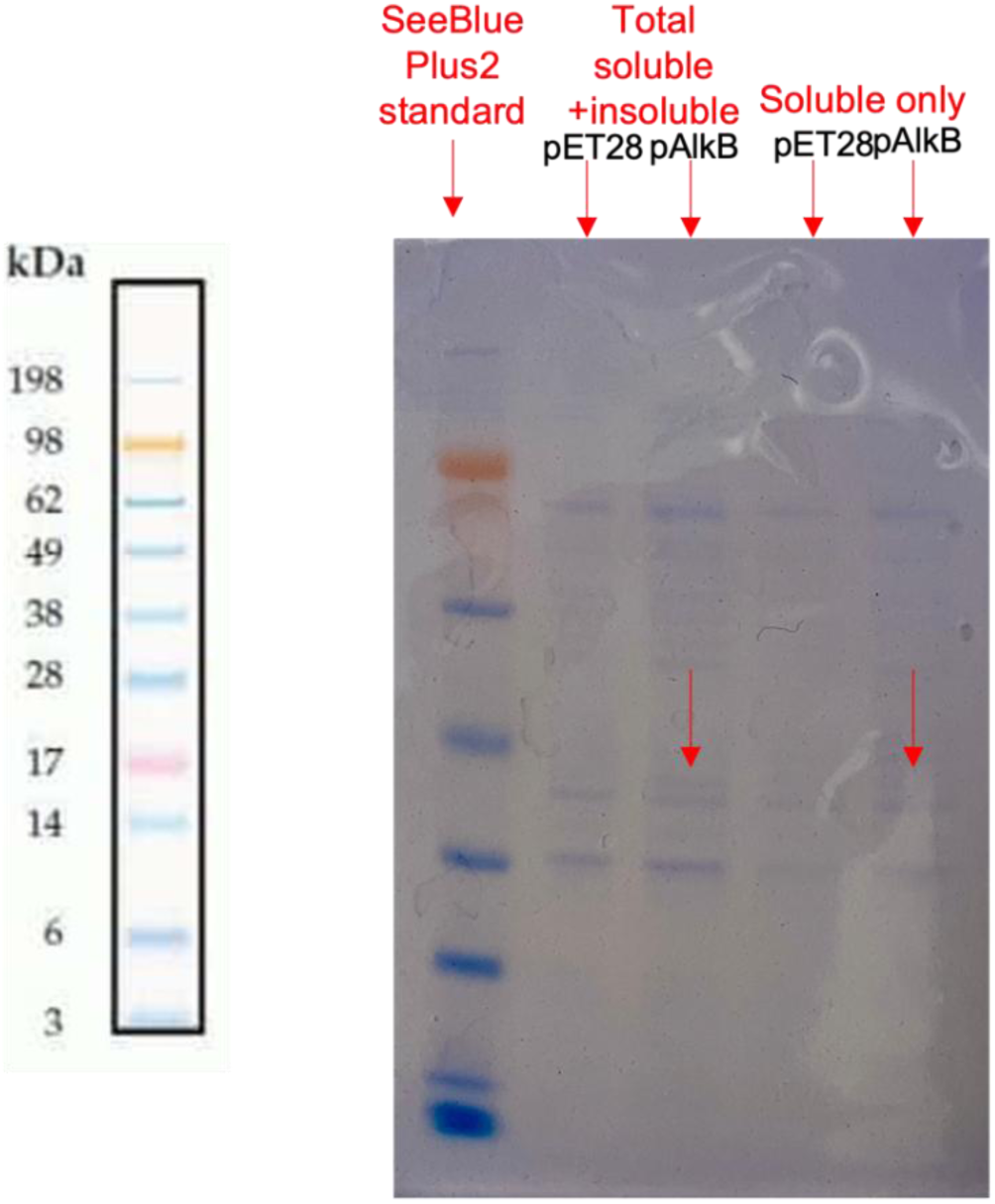
Polyacrylamide gel, showing a ∼46 kDa protein band (indicated with red arrows) in both the insoluble and soluble components. Standards from the SeeBlue Plus2 reference ladder are also included.

Protein expression was carried out at ambient temperature for 18 h, rather than the convention of 37 °C for 5 h. This was found to be a required optimization since no protein expression was detected by the former method. The rationale for this is two-fold; (i) it gives more time for a build-up of a higher concentration of protein, and (ii) lower temperatures (i.e., 15-20 °C) may facilitate more-precise protein folding and increase the protein stability [78].

## Acknowledgments

Special thanks go to Bethan J. Johnson (QUB) who provided useful discussion concerning the biofilm assays.

